# Dispersal, habitat filtering, and eco-evolutionary dynamics as drivers of local and global wetland viral biogeography

**DOI:** 10.1101/2023.04.28.538735

**Authors:** Anneliek M. ter Horst, Jane D. Fudyma, Jacqueline L. Sones, Joanne B. Emerson

## Abstract

Wetlands store 20-30% of the world’s soil carbon, and identifying the microbial controls on these carbon reserves is essential to predicting feedbacks to climate change. Although viral infections likely play important roles in wetland ecosystem dynamics, we lack a basic understanding of wetland viral ecology. Here 63 viral size-fraction metagenomes (viromes) and paired total metagenomes were generated from three time points in 2021 at seven fresh- and saltwater wetlands in the California Bodega Marine Reserve. We recovered 12,826 viral population genomic sequences (vOTUs), 4.4% of which were also detected at the same field site two years prior, indicating a small degree of population stability or recurrence. Viral communities differed most significantly across the seven wetland sites and were also structured by habitat (plant community composition and salinity). Read mapping to a new version of our reference database, PIGEONv2.0 (now with 515,763 vOTUs), revealed 196 vOTUs present over large geographic distances, often reflecting shared habitat characteristics. Wetland vOTU microdiversity was significantly lower locally than globally and lower within than between time points, indicating greater divergence with increasing spatiotemporal distance. Viruses tended to have broad predicted host ranges via CRISPR spacer linkages to metagenome-assembled genomes (whether this reflects true biology remains to be seen), and increased SNP frequencies in CRISPR-targeted major tail protein genes suggest viral eco-evolutionary dynamics, potentially in response to both immune targeting and to changes in host cell receptors involved in viral attachment. Together, these results highlight the importance of dispersal, environmental selection, and eco-evolutionary dynamics as drivers of local and global wetland viral biogeography.

## Introduction

Wetlands are an important carbon sink, estimated to store between 20-30% of the global soil carbon ^1^. They also provide ecosystem services, such as flood control, drought prevention, and water quality protection, and they support a rich biodiversity ^1–4^. However, these ecosystems are currently being lost at an estimated annual rate of 1.5% globally, releasing stored carbon into the atmosphere ^2, 5^. Moreover, due to climate change, soil salinity is increasing in formerly freshwater wetlands, causing changes to microbial and plant communities ^6–8^ and potentially leading to biodiversity loss ^9^. Microorganisms play central roles in carbon turnover and the emission of greenhouse gasses from wetland ecosystems^10^, and, by infecting, controlling the metabolism of, and lysing microorganisms, viruses also likely impact these biogeochemical cycles ^10, 11^. It is therefore important to characterize fresh- and saltwater wetland microbial and viral communities, in order to understand the ecological and biogeochemical responses of these fragile ecosystems under a changing climate ^12–14^.

While viruses are highly abundant in peat wetlands and other soils ^15–18^, we still know relatively little about wetland viral ecology, as methodological improvements have only recently made it possible to study soil viral communities in detail. While some prior efforts have focused on bioinformatic mining of viral sequences from total soil metagenomes ^19, 20^, by purifying the viral size fraction through 0.22 µm filtration prior to metagenomic sequencing (viromics), a much higher viral diversity can be recovered ^11, 15, 21^. Application of these methods to peatlands and a variety of other soils is beginning to reveal ecological factors important to soil viral biogeography.

Recent studies have shown substantial differences in soil viral community composition among habitats at both regional and global scales ^15, 22^. For example, soil viral ‘species’ (vOTUs) were rarely shared among four different habitats (grasslands, shrublands, woodlands, and wetlands) in northern California ^22^, and similarly, few RNA viral sequences were shared between grasslands and peatlands in the United Kingdom ^23^. Despite repeated observations of soil viral community heterogeneity at regional or continental scales ^18, 24^, the same viral ‘species’ (vOTUs) can be found on different continents, usually in the same habitat (*e.g*. peat viruses tend to be restricted to other peatlands) ^15^. While habitat seems to be an important contributor to soil viral biogeography, given the sparseness of the data, further studies are needed to assess the generalizability of these patterns.

At more local scales, soil viral community spatial structuring and temporal turnover have been observed, with viral communities showing seasonal dynamics ^25^ and exhibiting stronger spatial and temporal distance-decay relationships than bacterial communities ^18, 21^. However, those studies were conducted within the same habitat or soil type; differences in viral community composition across habitats have rarely been considered at local scales. In two studies that did compare viral communities by habitat in the same Swedish ecosystem, viral communities were found to be distinct among three habitats along a peatland permafrost thaw gradient ^16, 19^. However, those three habitats were also spatially separated, making the relative influences of habitat and spatial location difficult to disentangle. Similarly, viral community compositional differences along a grassland pH gradient also reflected spatial separation, but pH was seemingly the predominant factor driving viral community composition, which was corroborated in a meta-analysis of other soil and peat viral datasets ^26^. Disentangling the relative impacts of habitat characteristics and spatial location on soil viral community composition is thus an important near-term goal for advancing the field, but appropriate spatiotemporal scales for sampling soil viral communities are still unknown.

Building on our prior regional study of 30 viromes from four habitat types with very little overlap in viral ‘species’ (vOTUs) across samples ^22^, here we hypothesized that reducing complexity from the regional to local scale and restricting the diversity of habitats considered (only wetlands) would yield sufficient vOTU co-occurrence to link viral ecological patterns to their potential underlying drivers. We sampled seven different wetland sites across a 0.6 km² area at three time points in 2021 at the Bodega Marine Reserve on the California Pacific Coast (USA). We generated 63 viral size-fraction metagenomes (viromes) and 63 total soil metagenomes to profile the dsDNA viral communities and bacterial and archaeal (prokaryotic) communities, respectively, in these wetlands. We also compared results to our viromic dataset from Bodega Bay collected two years prior (in 2019) ^22^. Here we explore local and global wetland viral biogeography, investigate which factors among spatial distance, plant and microbial community composition, soil physicochemical properties, and time have the strongest influence on viral community composition, and evaluate the influences of spatial and temporal distance on viral population microheterogeneity and virus-host eco-evolutionary dynamics.

## Results and Discussion

### Dataset overview

To investigate wetland dsDNA viral biogeography on a local scale, we sampled seven nearby wetland sites within a 0.6 km² area in the Bodega Marine Reserve, California, USA (Figure 1A, map). Sampling sites were initially selected to represent freshwater, brackish, and saltwater wetlands, based on institutional knowledge of plant community composition, and we subsequently measured both plant communities and salinity to empirically define the sampled habitats. Near-surface (top 15 cm) wetland soils were collected at three time points (March, May, and July of 2021), with three replicate samples per time point per wetland site. Replicates were collected on average 17 m apart, with the closest samples within a site (regardless of the time point) 1.7 m apart and the farthest 89 m apart (Supplementary Table 1). All 63 samples (7 sites x 3 replicates x 3 time points) underwent viral size-fraction metagenomics (viromics) and total metagenomics to measure viral and prokaryotic community composition, respectively. A suite of soil physicochemical properties was also measured for each sample (Supplementary Table 2).

**Figure 1:**
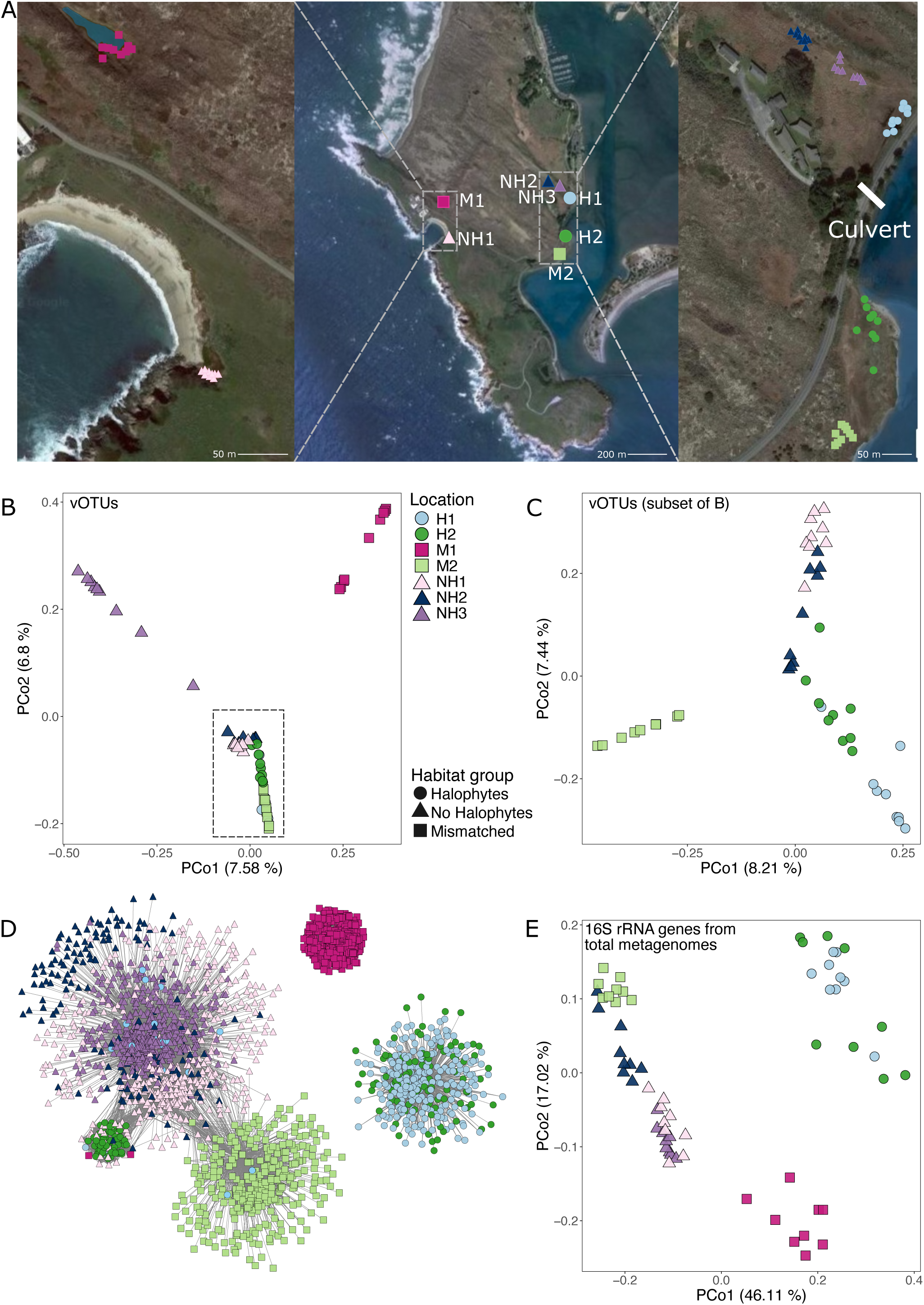
Sampling design and overarching compositional patterns for Bodega Bay viral and prokaryotic communities. A) Sampling locations for all Bodega Bay samples. Center: locations of the seven wetland sites within the Bodega Marine Reserve, Left and Right: locations of each of the nine samples per site (a zoomed in view of each site with individual sample labels is in Supplementary Figure 1). Per the legend below the images, circles correspond to locations with halophyte vegetation and saline soils, triangles correspond to locations without halophytes and non-saline soil, and squares correspond to mismatched locations. The ‘culvert’ label indicates the location of a human-made pipe below the road that allows for water movement. B-C) Principal coordinates analysis (PCoA), based on Bray-Curtis dissimilarities derived from the table of vOTU abundances (read mapping to vOTUs). Each point is one sample (one virome), with viral communities from B) all 63 viromes, and C) the 45 viromes indicated by the dashed rectangle in B. Panel C is a new PCoA to better show separation among overlapping samples in B. D) Co-occurrence network of vOTUs detected in more than one Bodega Bay virome, colored by the site in which they were most abundant (had the highest average per-bp coverage depth). Nodes represent vOTUs, and edges represent a significant co-occurrence between the vOTUs, calculated using a probabilistic co-occurrence model with the R package cooccur. E) PCoA based on Bray-Curtis dissimilarities of 16S rRNA gene OTU community composition from 63 total metagenomes. For all PCoA plots (B, C, E), the percent variance explained by each axis is indicated in parenthesis.

In total, 12,826 viral operational taxonomic units (vOTUs, ≥ 10 kbp, ≥ 95% average nucleotide identity, approximately species-level taxonomy ^27^) and 219 metagenome-assembled genomes (MAGs, ≥ 50% complete, ≤ 10% contaminated ^28^, Supplementary Table 3) were detected in our samples. From the viromes, we assembled 17,703 viral contigs *de novo*, which clustered into 12,261 vOTUs, and we recovered an additional 565 vOTUs by read mapping to our Phages and Integrated Genomes Encapsidated Or Not (PIGEONv2.0) database of 515,763 vOTUs from diverse ecosystems, including 369 vOTUs recovered from Bodega Bay viromes collected in 2019 ^22^. Read mapping to these sets of vOTUs and MAGs yielded the estimated relative abundances of each vOTU and MAG in each sample, used for downstream community compositional analyses (Supplementary Tables 4,5).

### Habitat features (plant community composition and salinity)

We identified 32 plant species across the seven sites (Supplementary Table 6), and plant communities separated the sites into two vegetation groups, based on the presence or absence of halophytes (salt-tolerant plants). There were no overlapping plant species between the two groups, and while most sites in the same vegetation group shared at least one dominant plant species, plant community composition differed at each of the seven sites (Supplementary Table 6). We also used salinity measurements to define habitats, with electrical conductivity measurements ranging from 0 to 82 mmhos/cm in our wetlands, and those between 0 to 2 mmhos/cm considered non-saline, 2 to 4 slightly saline, 4 to 8 moderately saline, 8 to 16 strongly saline, and 16 or greater extremely saline wetlands ^29^. Although vegetation tended to be indicative of soil salinity, our salinity measurements varied both within and among sites, and two sites had consistently mismatched salinity and vegetation measurements (site M1 had a ‘no halophyte’ plant community with non-to-strongly saline soils, and site M2 had a ‘halophyte’ plant community with non-saline soils). These seemingly contradictory vegetation and salinity results left us initially concerned that our salinity measurements might have been faulty, but evaporation during dry periods, seasonal waterlogging, precipitation, and leaching of water can all influence soil salinity in short time spans ^30^. Halophytic plants outcompete non-halophyte plants in saline environments ^31^, and coastal salt marshes such as site M2 experience tidal flooding with seawater, leading us to believe that M2 likely sometimes experiences higher soil salinity than we measured, promoting halophyte growth. Halophytes are not competitive in non-saline habitats ^32^, and since there were no halophytes at site M1 despite the moderate salinity measurements, we speculate that M1 soil is often non-saline. Regardless of the underlying mechanism(s) for the differences, we separated the seven sites into four habitat groups: “Halophyte (H)’’ for the two sites with halophyte plants and overall medium to extreme soil salinity (H1 and H2), “No Halophyte (NH)’’ for the three sites with no halophytes and low to slight soil salinity (NH1, NH2, and NH3), and two “Mismatched (M)” groups (M1 and M2) for the two sites for which the vegetation did not correspond with soil salinity. Importantly, the mismatched (M) sites did not share the same vegetation and salinity mismatch, so they do not represent the same habitat type, leading to four habitat groups (H, NH, M1, M2).

### Viral and prokaryotic communities were distinct at each of the seven wetland sites but were more similar within than between habitat types

Most (90%) of the viral ‘species’ (vOTUs) were restricted to only one of the seven wetland sites. While 38% of the vOTUs were detected in only one of the 63 viromes (Supplementary Figure 1), the proportion of these ‘singleton’ vOTUs was substantially reduced, compared to our prior regional-scale comparison of 30 viromes across grassland, shrubland, woodland, and wetland habitats, in which 81% of the vOTUs were detected in only one virome ^22^. Thus, the localized focus in one area and restriction to wetland habitats here, as well as increased spatiotemporal resolution, improved our ability to identify vOTUs shared across samples, as is necessary for recognizing biogeographical patterns. Of the 62% of vOTUs detected in more than one virome, 6,680 (52%) were recovered only within one wetland site, and viral community composition was significantly different at each site (PERMANOVA p < 0.001, Figure 1B, 1C). Viral community beta-diversity was significantly negatively correlated with spatial distance (Supplementary Figure 2A), implicating dispersal limitation as one potential driver of these patterns (as also suggested in previous work ^18, 21, 22^).

Despite overarching differences among wetland sites, viral communities grouped secondarily according to habitat type (Figure 1B, 1C), with significant differences among the four habitat groups (H, NH, M1, and M2, PERMANOVA, p < 0.001). Consistent with edaphic factors as potential drivers of these differences, viral community composition correlated significantly with soil chemical measurements (Supplementary Figure 2B, Supplementary Table 2), such as pH, moisture content, and sulfate concentrations (Supplementary Figure 2C). Considering only between-habitat beta-diversity, the viral communities from sites NH1 and NH2 were the most similar (Figure 1C), despite being physically far apart (Figure 1A), perhaps related to their similar salinity and plant communities (Supplementary Table 6). This is consistent with prior work that has suggested that plant cover type could play an important role in shaping soil viral communities ^33^. Similar salinity and plant communities were likely also drivers of viral community compositional similarity at sites H1 and H2. Those sites are also connected by a culvert (a human-made water tunnel beneath a road) (Figure 1A), presumably facilitating dispersal between the sites. Finally, the two mismatched sites each had distinct viral communities, potentially due to their unique combinations of plant composition and salinity.

A co-occurrence analysis revealed that vOTUs were most often shared across samples from the same habitat type. Specifically, vOTUs from all three non-halophyte soils (NH1, NH2, and NH3) tended to co-occur, as did vOTUs from wetlands with halophyte plants (H1 and H2). Perhaps reflecting the lack of other samples from the same habitat types in this dataset, vOTUs from each of the mismatched sites tended to co-occur only with other samples from the same site. Interestingly, a small subnetwork of vOTUs from the halophyte site H2 co-occurred with vOTUs from the non-halophyte wetlands. All of those co-occurring vOTUs were either the most abundant in or only detected in one particular H2 sample, H2-1-T2, which had low salinity (1.69 mmho/cm) (Supplementary Table 2). This suggests that environmental selection (presumably by way of microbial hosts) can act on wetland viral communities on very short time scales, and/or that an influx of new vOTUs was brought to site H2, being already adapted to conditions in the less saline water that brought them there.

The two sites with the most within-site vOTU co-occurrences also had the highest moisture content, consistent with hydrological mixing facilitating greater viral community homogeneity. Specifically, site NH3 harbored viral communities distinct from all other sites (Figure 1B), despite its similar salinity and plant community composition to the other two non-halophyte sites and its close proximity to NH2 (Figure 1A, Supplementary Table 2). A comparatively large percentage of vOTUs was shared across samples within the NH3 site (35% of vOTUs were shared among five or more NH3 samples, relative to only 8% on average for the other two non-halophyte sites, Supplementary Figure 1). Similarly, communities from site M1 were also distinct, with 30% of their vOTUs detected in five or more samples from the same site, whereas the five other sites (not NH3 or M1) shared only 9% of their vOTUs across five or more samples from the same site. Soil moisture content was highest at sites NH3 (83% on average) and M1 (52% on average), compared to 34% on average at the other five sites, likely facilitating more mixing and greater viral community homogeneity due to greater hydrologic connectivity. Overall, the viral community compositional and vOTU co-occurrence patterns revealed both dispersal (and dispersal limitation) and environmental selection (biotic and abiotic habitat characteristics) as likely drivers of local wetland viral biogeographic patterns.

To determine whether prokaryotic communities exhibited similar patterns to the viral communities, prokaryotic community composition and co-occurrence were also investigated. Briefly, the relative abundances and co-occurrences of MAGs and, separately, of 16S rRNA gene fragments recovered from total metagenomes were used for these analyses. While most of the prokaryotic communities were significantly different at each of the wetland sites (Figure 1E, Supplementary Figure 3A), the communities from the Halophyte sites (H1 and H2) were not significantly different from each other (PERMANOVA, p=0.055), grouping more by habitat type than did the viral communities. Co-occurrence networks for MAGs showed similar patterns to those of the viral communities, largely reflecting shared MAGs within the same habitat type, though relatively few MAGs were recovered from the non-halophyte wetlands (Supplementary Figure 3B,C). Although OTUs also showed the most co-occurrence within habitat types, OTUs were detected in multiple habitats far more often than were MAGs, suggesting that increased resolution (i.e., not requiring assembly into MAGs) revealed more co-occurrence, presumably due to increased access to rare community members. Overall, patterns for prokaryotic communities were similar to those of their viruses, and viral community composition was significantly correlated with prokaryotic community composition (Mantel test, p < 0.001), suggesting that at least some of the observed viral biogeographical patterns were due to habitat filtering (environmental selection) by way of their hosts.

### Global distribution patterns for Bodega Bay vOTUs suggest that wetland viral biogeography reflects habitat and salinity

To compare vOTUs recovered at Bodega Bay to the global viral metacommunity, we leveraged a new version of our Phages and Integrated Genomes Encapsidated Or Not (PIGEONv2.0) database, which we introduce here. Since the first iteration of PIGEON (PIGEONv1.0), which contained 266,125 vOTUs ^15^, PIGEONv2.0 has almost doubled in size, now including 515,763 vOTU sequences. Most notably, we increased the number of soil vOTUs from 15,892 to 61,757, predominantly from our in-house soil viromics data, including previously unpublished datasets that we are now making publicly available in PIGEONv2.0. The number of freshwater vOTUs also substantially increased, largely due to the addition of viruses from aquatic viromes from Lake Baikal in Russia ^34^. Here, these PIGEON improvements have facilitated global comparisons of Bodega Bay vOTU occurrence patterns.

Of the 12,826 vOTUs recovered at Bodega Bay, 196 (1.5%) were previously detected at other sites throughout the world (Figure 2A), recovered here through read mapping to PIGEONv2.0 (Figure 2B, Supplementary Table 7). Bodega Bay vOTUs were previously recovered from non-wetland soils (83), freshwater lakes (57), marine ecosystems (33), non-peat freshwater wetlands (14), and peat wetlands (8), indicating globally present viruses in relatively similar ecosystems throughout the world (Figure 2A, Supplementary Figure 4A). Notably, zero vOTUs from human-associated habitats were detected in these wetlands, perhaps indicating species boundaries between these very different habitat types. Most vOTUs that were detected in non-saline or slightly saline wetlands at Bodega Bay were originally recovered from non-wetland soils (62) or freshwater ecosystems (46), whereas most vOTUs from saline wetlands were previously recovered from marine (34) or non-wetland soil (28) ecosystems (Supplementary Figure 4B), again suggesting that habitat characteristics underlie global viral biogeographic patterns. Similarly, we also considered the relationship between vegetation group at Bodega Bay and the habitat in which a given vOTU was originally recovered (Figure 2D) and found that vOTUs from the non-halophyte sites were most often previously detected in non-wetland soils (79) or freshwater ecosystems (57), while vOTUs from the halophyte sites were most often previously detected in marine ecosystems (24) (Figure 2C). The detection of marine vOTUs in these wetlands is counter to our previous study of freshwater peatlands in Minnesota, USA, in which zero marine vOTUs from PIGEONv1.0 were detected ^15^, consistent with salinity as a habitat filter for vOTUs in both oceans and wetlands. Together, these results indicate that habitat characteristics – in this case, salinity and salinity indicators (halophyte or non-halophyte plant community composition) – can drive wetland viral community biogeography on a global scale.

**Figure 2:**
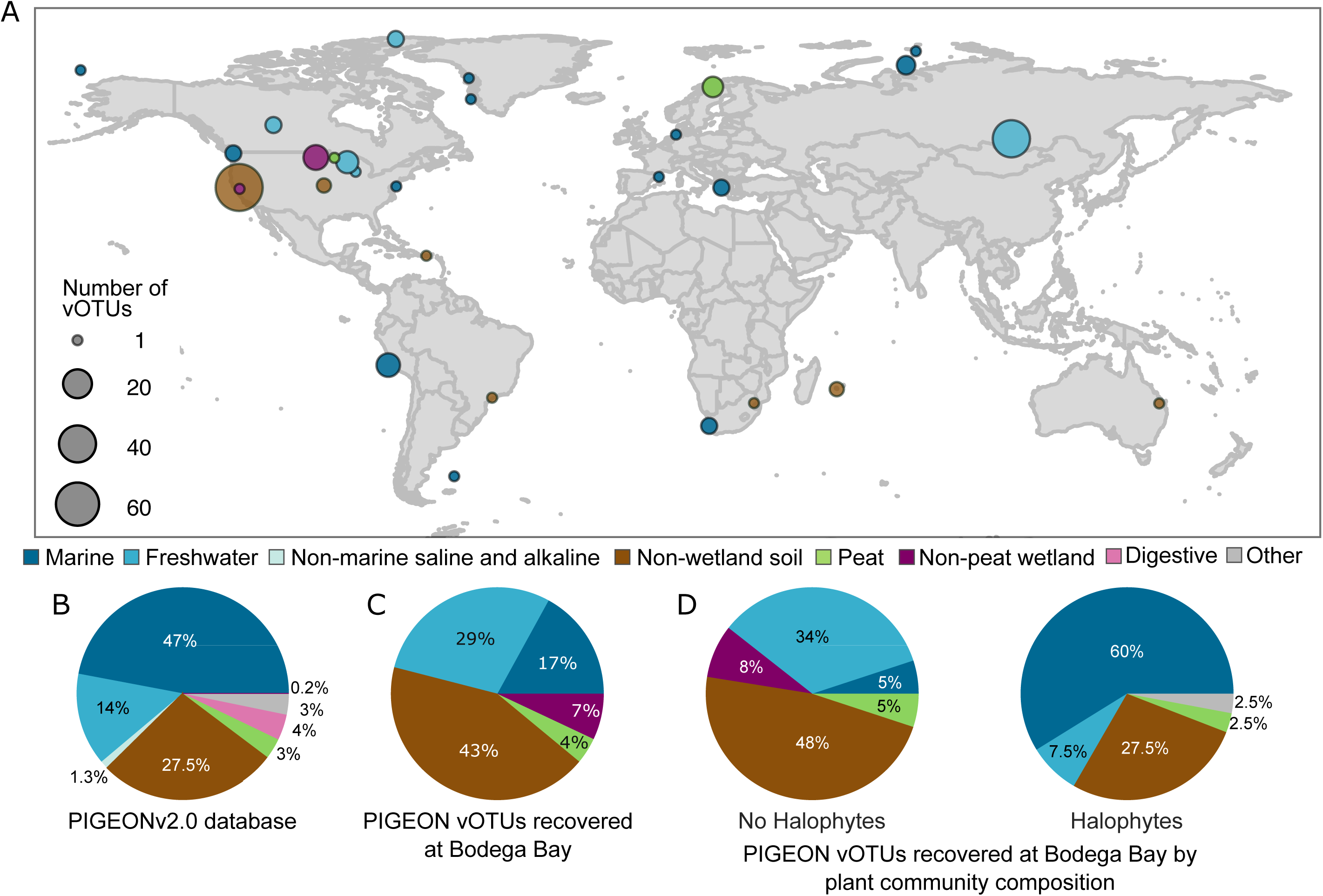
Global distribution and habitat context of Bodega Bay vOTUs, leveraging the PIGEONv2.0 database. A) vOTUs (n=196) from PIGEONv2.0 recovered at Bodega Bay by read mapping, according to the location where they were first recovered, colored by the environment in which they were originally recovered. Circle size indicates the number of vOTUs. B) Composition of the PIGEONv2.0 database of 515,763 vOTU sequences, colored by environment. C) Relative proportions of all vOTUs recovered from PIGEONv2.0 at Bodega Bay, colored by the original environment from which they were recovered. D) Relative proportions of vOTUs recovered from PIGEONv2.0 at Bodega Bay, as in panel C, but separated by the Bodega vegetation group in which they were recovered, colored by original source environment. If a vOTU was recovered in both vegetation groups, it appears twice in the chart.

### Wetland viral microdiversity was lower locally than globally and lower within than between time points

To investigate the contributions of viral genotypic heterogeneity to local and global viral ecology, we used inStrain ^35^ to calculate vOTU microdiversity profiles and compared dominant allelic variants over time and space. Specifically, we compared vOTU reference sequences initially recovered from PIGEONv2.0 (not assembled from Bodega Bay, 196), assembled from Bodega Bay in 2019 (2,377) ^22^, and assembled from Bodega Bay in 2021 (this study, 12,261) to their variants recovered in different samples at Bodega Bay. For each vOTU, we calculated pairwise average nucleotide identities (ANIs) between each sample-specific consensus variant sequence from Bodega Bay and the reference vOTU sequence. Genomic similarity between Bodega Bay variants and PIGEON references was significantly lower on average (average ANI 97.48%) than that for variants that were both assembled and recovered from Bodega Bay (average ANI 99.55%, Figure 3A, p <0.001, Student’s T-test). Given the global scale of PIGEON and local scale of Bodega Bay, this indicates greater viral population allelic variance (genomic heterogeneity) with increasing distance and/or time between samplings, a pattern known as ‘isolation by distance’ that has been studied for geographic distance, whereby populations in closer proximity are more genetically similar than populations that are farther away ^36^.

**Figure 3:**
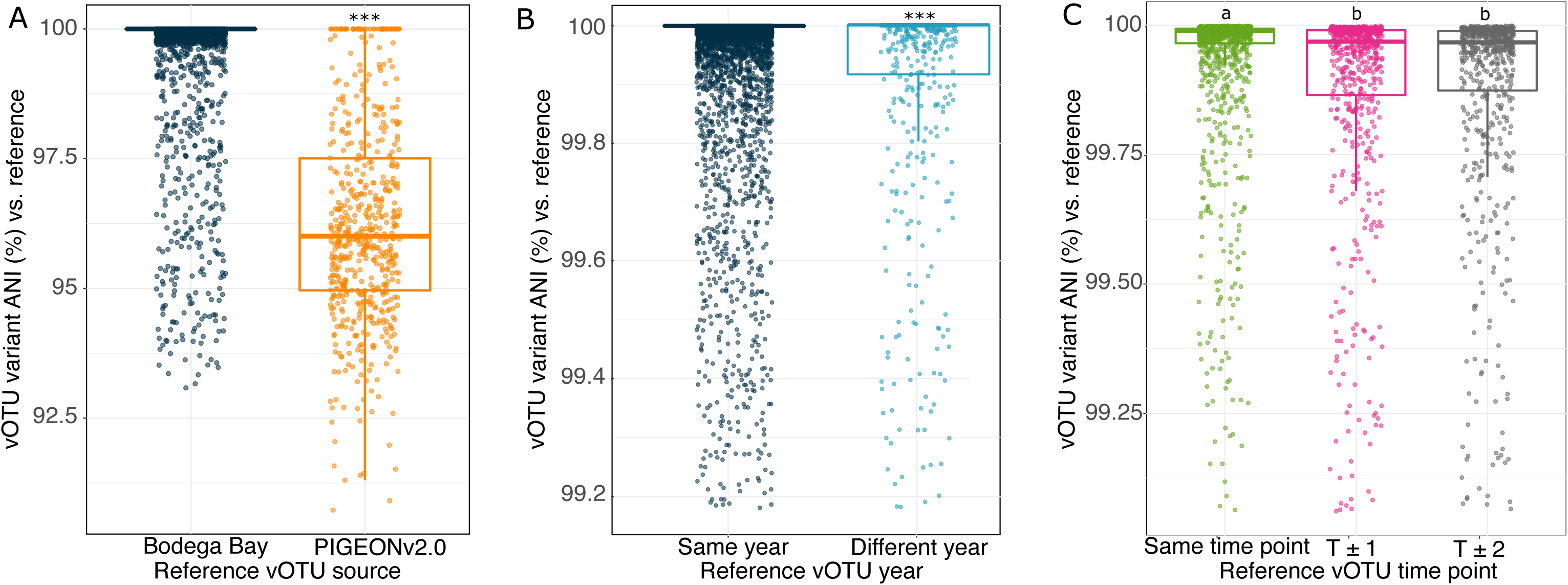
Comparisons of viral variant (sub-population) diversity in local and global contexts. Pairwise average nucleotide identities (ANIs) between vOTU variants, calculated between each sample-specific vOTU consensus sequence and the originally assembled (reference) vOTU sequence, using inStrain. Each point is the ANI for one vOTU variant in one Bodega Bay virome compared to the reference sequence for that vOTU. A) Variant ANIs for: (left) vOTUs both assembled and recovered through read mapping from the Bodega Bay dataset (Bodega Bay reference sequences), and (right) vOTUs recovered at Bodega Bay via read mapping but originally derived from PIGEONv2.0 (PIGEONv2.0 reference sequences). Stars above boxes correspond to significant differences between groups (Student’s T test, significant when p < 0.0001). B) Variant ANIs for vOTUs both assembled and recovered via read mapping from Bodega Bay, either: (left) assembled and recovered in the same year (2019-2019 or 2021-2021), or (right) in different years (2019-2021 or 2021-2019). C) Variant ANIs for vOTUs assembled from Bodega Bay in 2021, either assembled and recovered through read mapping at the same sampling time point, or at different time points, where T±1 equals 2 months between samplings, and T±2 equals 4 months. Letters above boxes correspond to significant differences between groups (Student’s T test, significant when p < 0.0001). In all three panels, boxes show the median and interquartile range (IQR), and whiskers extend to Q1-1.5*IQR and Q3+1.5*IQR.

A relatively small number of the Bodega Bay vOTUs detected in 2021 were also recovered from Bodega Bay in 2019 (568 vOTUs, 4.4% of the 2021 dataset). This suggests that a small part of the wetland soil virosphere was stable or consistently recurrent over time. However, for vOTUs that were assembled and recovered through read mapping in the same year, the genomic similarity of dominant allelic variants was higher (99.86%) than for vOTUs that were assembled and recovered in different years (99.25%, Student’s T-test, p<0.001, Figure 3B). Thus, although these viral ‘species’ persisted over time, their strain-level heterogeneity increased over the two years, consistent with temporal ‘isolation by distance’ ^36^, with populations farther apart in time exhibiting more genomic divergence.

Sub-population dynamics for vOTUs that were recovered multiple times within the same Bodega Bay wetland site in 2021 were also compared to assess short-term eco-evolutionary dynamics. Genomic similarity of dominant allelic variants was highest for vOTUs recovered through read mapping at the time point from which they were assembled (Figure 3C) and was significantly lower at both of the other time points. This indicates that, even over short time scales of one to two months, variants significantly fluctuated in abundance and/or diverged. Given that there was no linear trajectory in variant ANI divergence with time (variants were just as different between adjacent time points as between the first and third time points), abundance fluctuations seem more likely to explain these patterns than divergence.

We also used inStrain to compare MAG allelic variants in the 2021 Bodega Bay dataset. MAG variants recovered and assembled at the same time point were most genomically similar (had the highest ANI), whereas MAGs from different time points had significantly lower ANI (Supplementary Figure 3D). Interestingly, in contrast to the vOTU variants, MAG sub-population dynamics exhibited temporal progression, with sub-population pairs from the same time point most similar, those from the first and last time points most distinct, and those from adjacent time points (i.e., from time points 1 and 2 or 2 and 3) exhibiting intermediate similarity in their ANIs. Additional time points would be required to determine whether this is likely due to divergence over time, but results show sub-population dynamics for both viral and prokaryotic populations over months.

### Viral ‘species’ (vOTUs) tended to have broad predicted host ranges, and on average, MAGs had evidence for interactions with more than 10 vOTUs past

To investigate putative host ranges, we bioinformatically linked vOTUs to MAGs, using CRISPR arrays ^37^. All 12,826 vOTUs and 219 MAGs (210 Bacteria and 9 Archaea) were used for this analysis. A total of 29,709 CRISPR arrays was recovered from the metagenomes, and 683 virus-host linkages were predicted between 378 vOTUs and 53 MAGs. All identified host MAGs were bacteria and could be classified to at least the phylum level, with Proteobacteria and Actinobacteriota among the most commonly reconstructed MAGs (Figure 4A). Samples from medium to extremely saline wetlands had significantly more CRISPR arrays and spacers than others, perhaps suggesting increased viral predation, but there was no significant relationship between the number of CRISPR arrays or spacers and the number of vOTUs in a given sample (Supplementary Figure 5). The average MAG was linked to 13 vOTUs, indicating that wetland prokaryotic populations can be infected by (or otherwise interact with ^38^) multiple, diverse viral species. On average, each vOTU was linked to four MAGs, and 164 vOTUs (45% of those with predicted hosts) had putative linkages to MAGs in different phyla (Figure 4A, 4B). The average vOTU was linked to MAGs in two phyla, and when only considering vOTUs linked to more than one MAG, vOTUs were linked to MAGs across three or more phyla on average.

**Figure 4:**
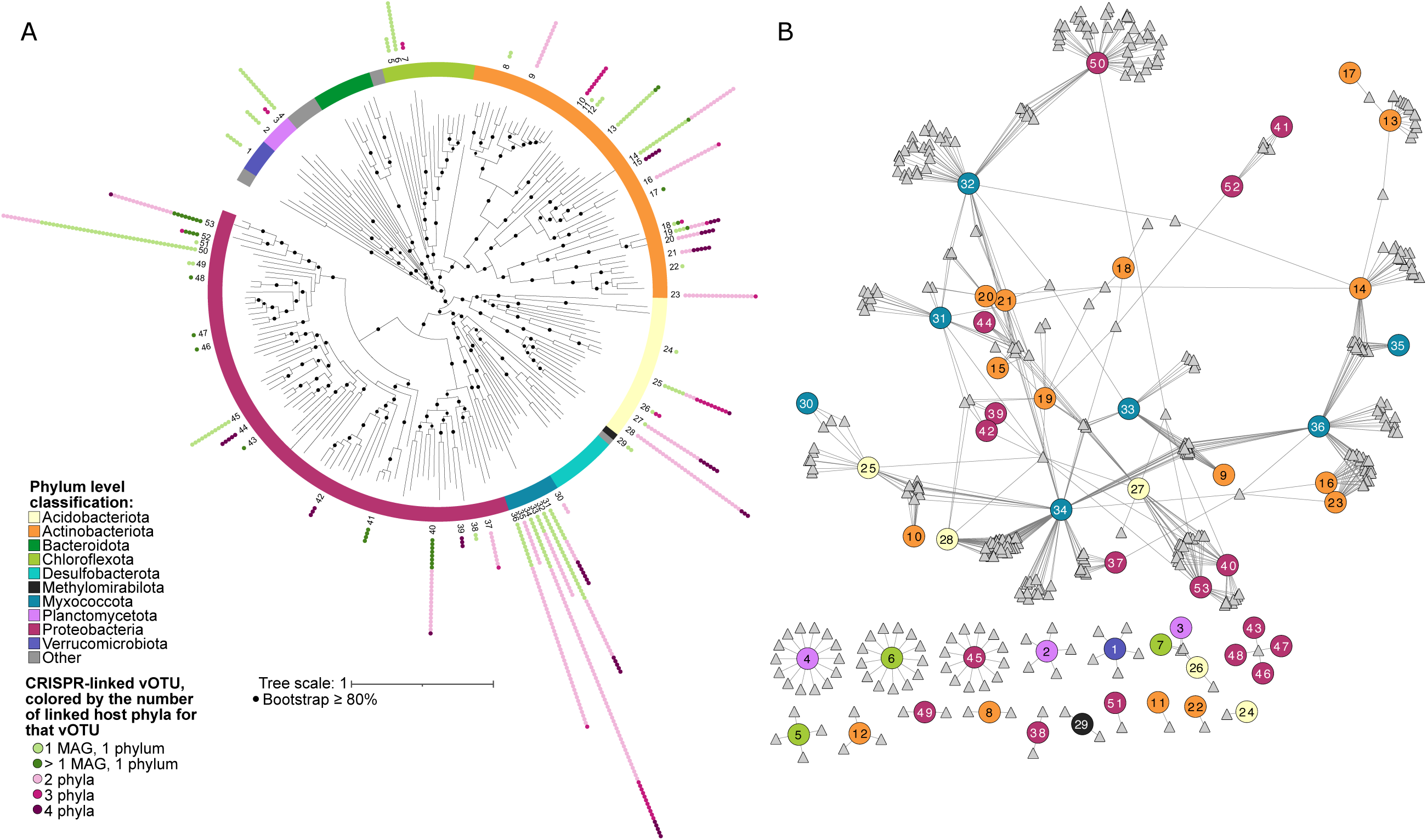
Bodega Bay virus-host linkages and putative interactions derived fromCRISPR spacer-protospacer matches. A) Unrooted phylogenetic tree (concatenated predicted protein alignment of 43 marker genes defined by CheckM) of prokaryotic metagenome-assembled genomes (MAGs) with at least one vOTU linked by CRISPR sequence homology. The numbers for MAGs correspond to numbers in the network in panel B. Tree was constructed using gtdbtk under the WAG model. B) Virus-host linkage network for MAGs with at least one vOTU linked through CRISPR homology. Circle nodes represent MAGs and are colored by phylum, while triangles represent vOTUs.

These results suggest either that CRISPR spacer matches to viral proto-spacers are imperfect for predicting virus-host linkages associated with infections in these systems, or that wetland viruses have much broader host ranges than previously appreciated. Recent studies have suggested that viral interactions with hosts may be far less specific than previously understood, with viruses infecting (or otherwise interacting with) prokaryotes across different phyla ^38, 39^. The mechanisms that could routinely enable viruses to infect different phyla are unknown, but recent evidence for diverse plasmid-dependent phages ^40^ (which target conjugation proteins encoded by horizontally transferrable plasmids) offers one interesting possibility for cross-infection that bears further exploration. Cross-phylum CRISPR linkages could also reflect non-specific interactions (e.g., uptake of viral particles or DNA by non-primary hosts, or horizontal transfer of CRISPR regions), as opposed to infections, and these interactions have been suggested to be more common than previously appreciated ^38^.

To investigate viral evolution in response to host immunity, we calculated the allelic variance within and outside of the viral genomic regions linked to CRISPR spacers, using the originally assembled vOTU sequence as the reference for SNP identification for each vOTU. Viral genomic regions with a CRISPR-spacer match had on average 5.6 SNPs/Kbp, whereas the genome outside of the match had on average 3.3 SNPs/Kbp, indicating more allelic variance in CRISPR-targeted regions, compared to the rest of the viral genome. This has been seen previously, for example in *Streptococcus thermophilus* phage-host coevolution experiments and in an acid mine drainage system^41, 42^, and it suggests increased phage genome diversification in CRISPR targeted regions to promote immune evasion. Of the predicted proteins in the CRISPR-targeted viral genomic regions with SNPs, 87% were annotated as hypothetical proteins, and 9% were putative major tail proteins. A significantly larger proportion of putative tail proteins were found in these regions than were annotated as putative major tail proteins in the whole dataset (0.93%, p < 0.00001, Z-test). This suggests that there is selection for accelerated evolution in viral genomic regions targeted by CRISPRs, particularly in tail proteins likely involved in attachment to host cell receptors ^43^. Evidence for higher mutation rates in phage tail protein genes is presumably due to viral adaptation to changes in host cell receptors to facilitate attachment, as previously suggested ^44, 45^.

## Conclusions

Here, we analyzed dsDNA viral communities from the Bodega Bay, California wetland ecosystem and showed significant differences in viral community composition across seven wetland sites, with evidence for dispersal, dispersal limitation, and habitat filtering as underlying drivers of the observed patterns. Although wetland viral communities differed predominantly by location within Bodega Bay, perhaps reflecting local dispersal limitation, the two wetland sites with the most homogeneous communities had the highest soil moisture content, suggesting hydrologic mixing and more opportunities for within-site dispersal with increasing moisture content. Local wetland viral communities were secondarily structured by habitat characteristics, such as plant community composition and soil salinity, indicating environmental filtering, perhaps by way of host adaptation. A small fraction (1.5%) of the vOTUs were previously recovered elsewhere, with global biogeographical patterns largely linked to habitat characteristics; marine vOTUs tended to be recovered in saline wetlands, freshwater vOTUs in non-saline wetlands, and soil vOTUs across wetland habitats.

In addition to dispersal and environmental filtering, eco-evolutionary dynamics (e.g., diversification and/or compositional shifts among dominant allelic variants) contributed to local and global viral biogeographical patterns. Pairwise ANI % between dominant allelic variants (sub-populations) differed significantly between years and over the four-month timescale of this study. In addition, Bodega Bay vOTU variants tended to be more divergent from reference vOTUs recovered elsewhere globally than from reference sequences assembled from Bodega Bay. The observed greater divergence over larger spatiotemporal scales is consistent with patterns of ‘isolation by distance’, whereby variants closer together in time and/or space likely had greater opportunities for gene flow. On a global scale, this may reflect local diversification and global dispersal limitation of most variants. Our limited ability to link viruses to their hosts (a limitation of the current state of the field) makes the contributions of virus-host co-evolutionary dynamics to biogeographic patterns difficult to evaluate, but we did see evidence for virus-host interactions spanning multiple phyla. Taken together, these results highlight dispersal, environmental filtering, and eco-evolutionary dynamics as likely drivers of both local and global wetland viral biogeographical patterns, expanding our understanding of the highly diverse and dynamic global soil virosphere.

## Acknowledgements

We thank Sara Geonczy and Devyn Durham for their help with sampling, Christian Santos-Medellín for helpful discussions, and the University of California, Davis Natural Reserves site directors and staff, particularly Suzanne Olyarnik at the Bodega Marine Reserve, for facilitating site access and providing logistical support. Funding for this work was provided by the U.S. Department of Energy (DOE), Office of Science, Office of Biological and Environmental Research (BER), Genomic Science Program, award number DE-SC0021198 (grant to JBE).

## Data Availability Statement

Raw sequencing reads are available at NCBI under BioProject number PRJNA913601, and sequence processing and statistical analysis code can be found on GitHub (https://github.com/AnneliektH/BodegaBay2021). The PIGEONv2.0 database vOTU sequences, the vOTU sequences recovered in this dataset, and MAGs recovered in this dataset are available on Dryad (https://datadryad.org/), using the following DOI: doi.org/10.25338/B8C934.

## Methods

### Field site and sample collection

Samples were collected three times over six months at the University of California, Davis Bodega Marine Reserve, in seven wetland soil ecosystems within the reserve (Supplementary Table 2). Sample collections were performed on March 17th, May 13th,and July 15th, 2021 (T1, T2 and T3, respectively) from each of seven distinct wetland sites.The plant community at each site was used as an indicator for soil salinity (Supplementary Table 6), such that the seven wetland sites were initially selected to represent three low-salinity and four high-salinity habitats, but subsequent analyses revealed more nuance in these habitat types (see main text). At each time point, three replicate surface soil samples (0-15 cm deep, 2.5 x 2.5 cm square area) were collected per wetland site, using a soil knife. The soil within each sample was homogenized and stored at −80 °C until further processing.

### Virome DNA extraction, library construction, and sequencing

Soil virions were enriched using a modified version of a previously published protocol ^46^. For each sample, 10 grams of soil were suspended in 30 mL of protein-supplemented phosphate-buffered saline solution (PPBS: 2% bovine serum albumin, 10% phosphate-buffered saline, 1% potassium citrate, and 150 mM MgSO_4_ in ultrapure water), briefly vortexed, placed on an orbital shaker (30 min, 400 rpm, 4 °C), and then centrifuged (10 min, 3,095 x g, 4 °C). Supernatant was then centrifuged twice (8 min, 10,000 x g, 4 °C) to remove residual soil particles. The purified supernatants were then filtered through a 0.22 µm polyethersulfone membrane to remove most cells. The resulting filtrate was ultracentrifuged (2 hrs 25 min, 32,000 rpm, 4 °C) to pellet the virions, using an Optima LE-80K ultracentrifuge with a 50.2 Ti rotor (Beckman-Coulter Life Sciences). Supernatants were decanted, and pellets were resuspended in 100 µl of ultrapure water. DNase treatment was not performed, as soil samples were stored frozen prior to processing, due to COVID-19 lockdown restrictions, and avoiding DNase treatment on such samples has been shown to improve viromic DNA yields without substantially compromising the viral ‘signal’ in the data ^47^. DNA was extracted from the viral-enriched fraction, using the DNeasy PowerSoil Pro kit (Qiagen, Hilden, Germany), following the manufacturer’s instructions, with an added step of a 10-min incubation at 65 °C before the bead-beating step. Libraries were constructed by the UC Davis DNA Technologies Core, using the DNA Hyper Prep library kit (Kapa Biosystems-Roche, Basel, Switzerland), and paired-end 150 bp sequencing was done using the NovaSeq S4 platform (Illumina) to an approximate depth of 10 Gbp per virome.

### Total DNA extraction, library construction, and sequencing

Total DNA was extracted from 0.25 g of soil per sample with the DNeasy PowerSoil Pro kit (Qiagen, Hilden, Germany), following the manufacturer’s instructions, with an added step of a 10-min incubation at 65 °C before the bead-beating step. Libraries were constructed by the UC Davis DNA Technologies Core, using the DNA Hyper Prep library kit (Kapa Biosystems-Roche, Basel, Switzerland), and paired-end 150 bp sequencing was done using the NovaSeq S4 platform (Illumina) to approximate depth of 20 Gbp per total metagenome.

### Soil chemistry and moisture

Soil moisture was defined by calculating the gravimetric water content of the soil. Soil chemistry measurements were performed by Ward Laboratories (Kearney, NE, USA). Briefly, soil pH and soluble salts were measured using a 1:1 soil:water suspension. Soil organic matter was calculated as percent mass loss on ignition. Nitrate was measured using a KCl extraction. Potassium, calcium, magnesium and sodium were measured using an ammonium acetate extraction. Zinc, iron, manganese and copper were measured using a DTPA extraction. Phosphorus was measured using the Olsen method and sulfate was measured using a Mehlich-3 extraction.

### Virome bioinformatic processing

Reads were trimmed using Trimmomatic v0.39 ^48^ to remove Illumina adapters and for quality trimming, using paired-end trimming, a sliding window size of 3:40, and a minimum read length of 50 bp. PhiX sequences were removed using BBDuk, from the BBMap v38-72 package ^49^, using k=31 and hdist=1. *De novo* assemblies were generated separately for each virome from the quality-trimmed, phiX-free reads, using MEGAHIT v1.0.6 ^50^, with k-min of 27, presets meta-large, and a minimum contig length of 1000 bp. Contigs were renamed, using the rename command from the BBMap package ^49^, using standard settings, and only contigs 10kbp were retained, using reformat from BBmap with the setting minlength=10000. Viral contigs were predicted using VIBRANT v1.2.0 ^51^, in virome mode and retained for downstream analyses if VIBRANT classified the contig as viral. Viral contigs were dereplicated into vOTUs using dRep v3.2.0 ^52^ at 95% ANI with a minimum coverage threshold of 85%, using the ANImf algorithm. Reads were mapped to the vOTUs, using Bowtie2 v2.4.2 ^53^ in sensitive mode, and the resulting samfiles were converted to bamfiles via SAMtools v1.15.1 ^54^. A coverage Table was produced using CoverM v0.6.1 ^55^, using CoverM contig with the mean coverage and a minimum covered fraction (breadth) of 75% (Supplementary Table 4). Reads were subsequently mapped back to our PIGEONv2.0 database, using CoverM with the same settings.

### Total metagenome bioinformatic processing

Read trimming, PhiX removal, and assembly were done the same way as for the viromes. Contigs were renamed using using the rename command from the BBMap package ^49^, using standard settings, and only contigs 2 kbp were retained, using reformat from BBmap with the settings minlength=2000. CD-hit v2007-013 ^56^ was used to deduplicate the contigs at approximately 99% ANI, using the -c 0.99 -aS 0.99 settings. Bowtie2 v2.4.2 ^53^ was used to map reads to the contigs, using sensitive mode, and the samfiles were converted to bamfiles using SAMtools v1.15.1 ^54^. A depth file for binning was created using MetaBAT2 v2.12.1 ^57^, using jgi_summarize_bam_contig_depths. Bins were then created using MetaBAT2 v2.12.1, using standard settings. dRep v3.2.0 ^52^ was used to dereplicate the bins, using primary clustering at 90%, secondary clustering at 95%, coverage method larger, a contamination threshold of 5%, and a coverage threshold of 30% ^58^. CheckM v1.0.13 ^59^ was used to estimate completeness and contamination of genome bins, and bins 50% complete and 10% contaminated were retained ^28^. RefineM v0.1.2 ^60^ was used to refine the recovered bins, using standard settings. Reads were mapped to these bins (metagenome-assembled genomes, MAGs) using Bowtie2 v2.4.2 ^53^ with default settings except setting --min-covered-fraction to 0.5 ^61^ (Supplementary Table 5). Phylogenetic trees were constructed using gtdbtk v2.1.0 ^62^, using the classify-wf command for phylogenetic inference and for aligning the identified marker genes. After this, gtdbtk infer was used to create the phylogenetic tree, using standard settings.

### Recovery and analysis of 16S rRNA gene sequences from metagenomes

SortMeRNA v4.2.0 ^68^ was used against the bacterial and archaeal SILVA databases ^69^ to recover reads containing 16S rRNA gene sequences from the total soil metagenomes. RDP tools v11^70^ was used to taxonomically classify the sequences, using the RDP database v18 as a reference ^71^. A count Table of the 16S rRNA gene OTUs was generated using the hier2phyloseq() function from the RDPutils package ^72^.

### PIGEONv2.0

To build further upon PIGEON 1.0 ^15^, we added more viral sequences mostly from in-house soil data, both published ^18, 21, 22, 47^ and currently unpublished, and from recent publications of viral ecology in soil ^26^, lakes ^34, 63^ and oceans ^64, 65^. We also mined a total soil metagenome dataset for viral sequences ^66^. A prefix was added to all sequence headers, in order to quickly identify what dataset the original sequence came from (Supplementary table 9). All sequences were dereplicated using cd-hit 2007-0131 ^56^, because the dataset was too large to use other programs for dereplication.

### Microdiversity profiles

Within-population genetic diversity was calculated using inStrain v1.4.0 ^35^. The bam files created by bowtie2 from the viromes were used as input for the inStrain profile option to identify divergent sites for each of the vOTUs. Variants were only called if they had a minimum coverage of 5 reads. MAG population genetic diversity was calculated the same way, using the bam files created by bowtie2 from the total metagenomes as input for nStrain.

### CRISPR-spacer analyses for virus-host linkages

Crass v1.0.1 ^37^ was used to assemble spacer and repeat sequences in the total metagenomes, using −l 4. All spacer sequences were then compared to the vOTUs, using blastN v2.7.1 ^67^, retaining hits with fewer than two mismatches and >95% nucleotide identity. All repeat sequences were compared to the MAGs using blastN, retaining hits that had no mismatches and 100% nucleotide identity (Supplementary Table 8).

### Data analysis and visualization

All statistical analyses were done using R v 4.1.0 ^73^. Analysis for viral community composition were done on the mean coverage vOTU abundance Table, unless otherwise noted. Bray-Curtis dissimilarities were calculated on log-transformed relative abundances, using the vegdist function from the vegan package v2.6-2 ^74^. PERMANOVA analyses were done using the adonis2 function from vegan. Principal coordinates analyses were performed with the pcoa() function from ape v5.4-2 ^75^. The BIO-ENV analysis was done using the bioenv function from vegan. Co-occurrence analyses for vOTUs, MAGs and 16S rRNA OTUs were done using the coocur package in R ^76^, using a presence-absence version of the abundance Tables. Only significantly positive co-occurrences (p<0.001) were used for visualization. Co-occurrence networks were visualized using Cytoscape v3.7.1 ^77^, using the edge-weighted spring embedded model, placing vOTUs that co-occur more frequently in closer proximity to each other in the figure. Upset plots were created using the UpSetR package v1.4.0 ^78^, using a presence-absence version of the vOTU abundance Table. All maps were created using the R package ggmaps ^79^. Pie charts and bar charts were created with Python v3.8, using matplotlib v3.4.2 ^80^ and seaborn v0.11.2 ^81^. The phylogenetic tree was created using the iTOL website ^82^, and the CRISPR-repeat network linking viruses to hosts was created using Cytoscape. All other plots were created, using the R package ggplot2 v3.3.5 ^83^. Correlation tests between community Jaccard Dissimilarity and spatial or environmental distance were done using the cor.test() function, using the pearson method with the alternative parameter set to two-sided. The linear regression slopes were calculated using the lm function, as has been done previously ^18^. All scripts are available at https://github.com/AnneliektH/BodegaBay2021.

