## Supplementary figures 1-5 for "Dispersal, habitat filtering, and eco-evolutionary dynamics as drivers of local and global wetland viral biogeography"

**Summary: This supplementary information contains supplementary figures 1 – 5**

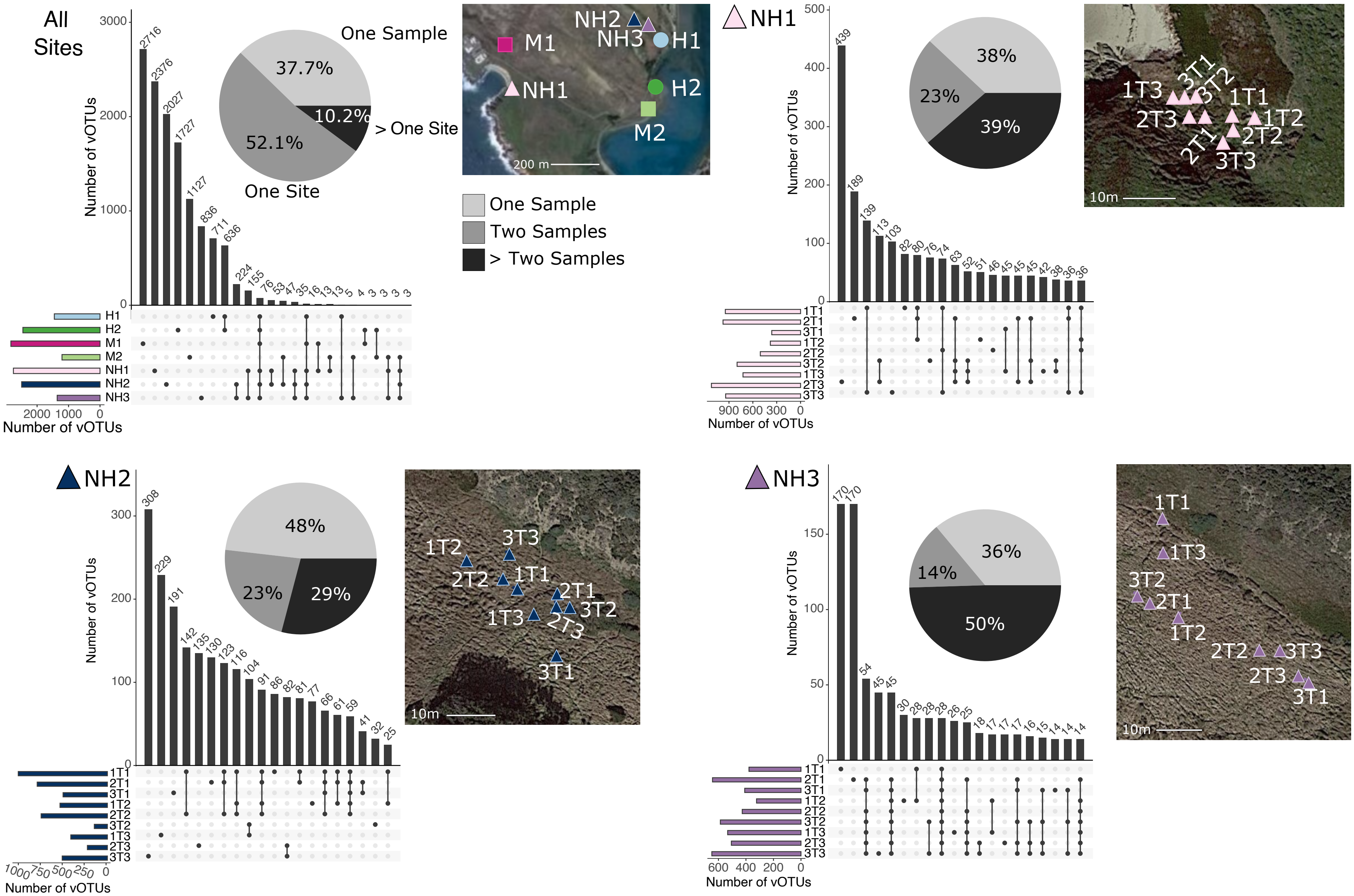


**Supplementary Figure 1 (page 1 of 2)**: UpSet plots of vOTU detection patterns (one for all sites together and one for each of the seven sites separately) with sampling locations and pie charts showing the percentage of vOTUs with a given detection pattern. For ‘all sites’, the detection patterns are: one sample only (light grey), one site (two or more samples within that site, dark gray), or two or more sites (black). For each wetland site, the detection patterns are: one sample (light grey), two or more samples from that site (dark gray), and three or more samples from that site (black). For each wetland site, detailed sampling locations are depicted, labeled by biological replicate number (1, 2, or 3 at the start of the label, indicating different locations at the same time point) and time point (T1, T2, or T3 at the end of the label). In the UpSet plots, black points indicate samples in which the vOTUs in the bar graph above were detected. The bars above each set of points indicate the number of vOTUs with that detection pattern.

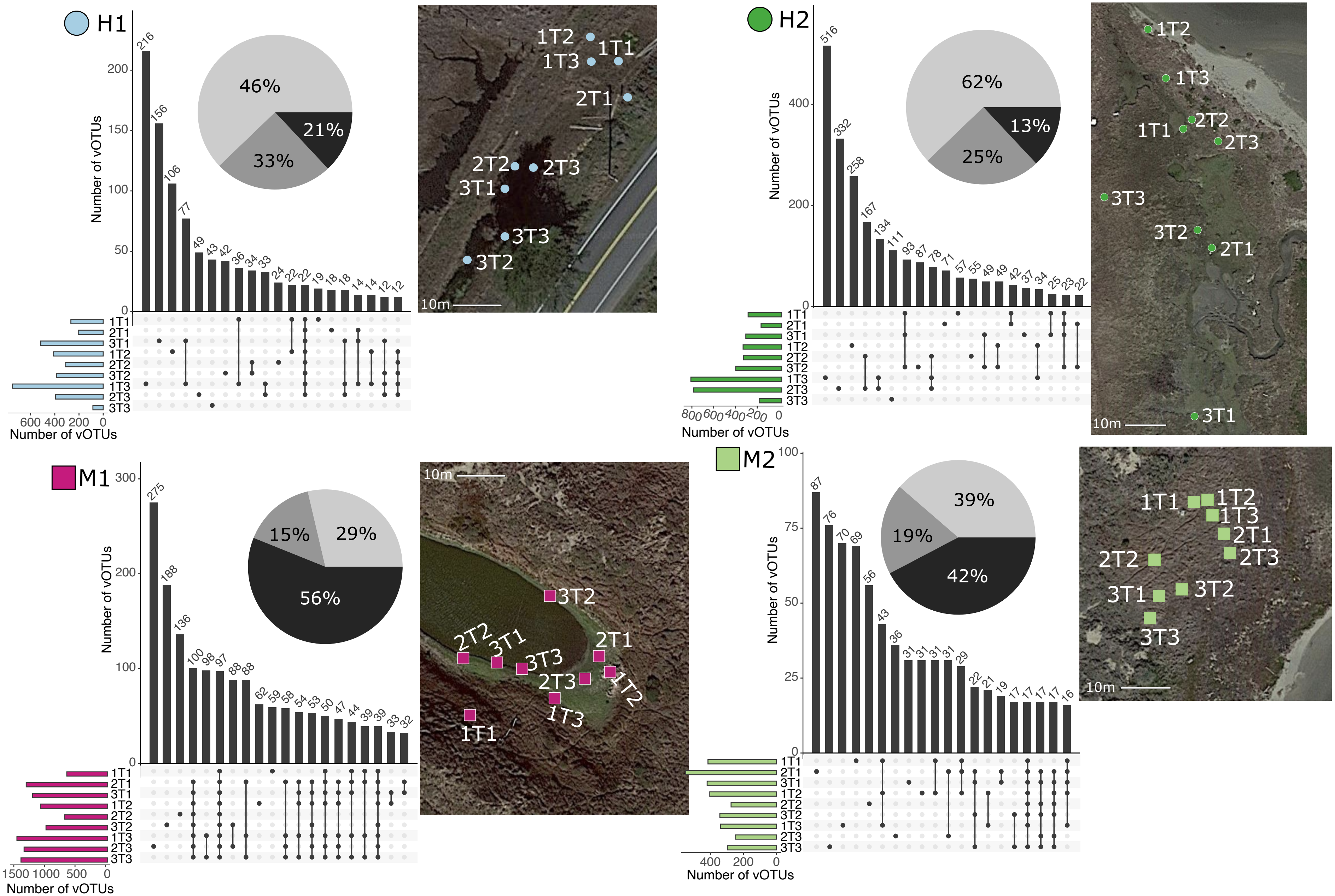


**Supplementary Figure 1 (page 2 of 2)**: UpSet plots of vOTU detection patterns (one for all sites together and one for each of the seven sites separately) with sampling locations and pie charts showing the percentage of vOTUs with a given detection pattern. For 'all sites', the detection patterns are: one sample only (light grey), one site (two or more samples within that site, dark gray), or two or more sites (black). For each wetland site, the detection patterns are: one sample (light grey), two or more samples from that site (dark gray), and three or more samples from that site (black). For each wetland site, detailed sampling locations are depicted, labeled by biological replicate number (1, 2, or 3 at the start of the label, indicating different locations at the same time point) and time point (T1, T2, or T3 at the end of the label). In the UpSet plots, black points indicate samples in which the vOTUs in the bar graph above were detected. The bars above each set of points indicate the number of vOTUs with that detection pattern.

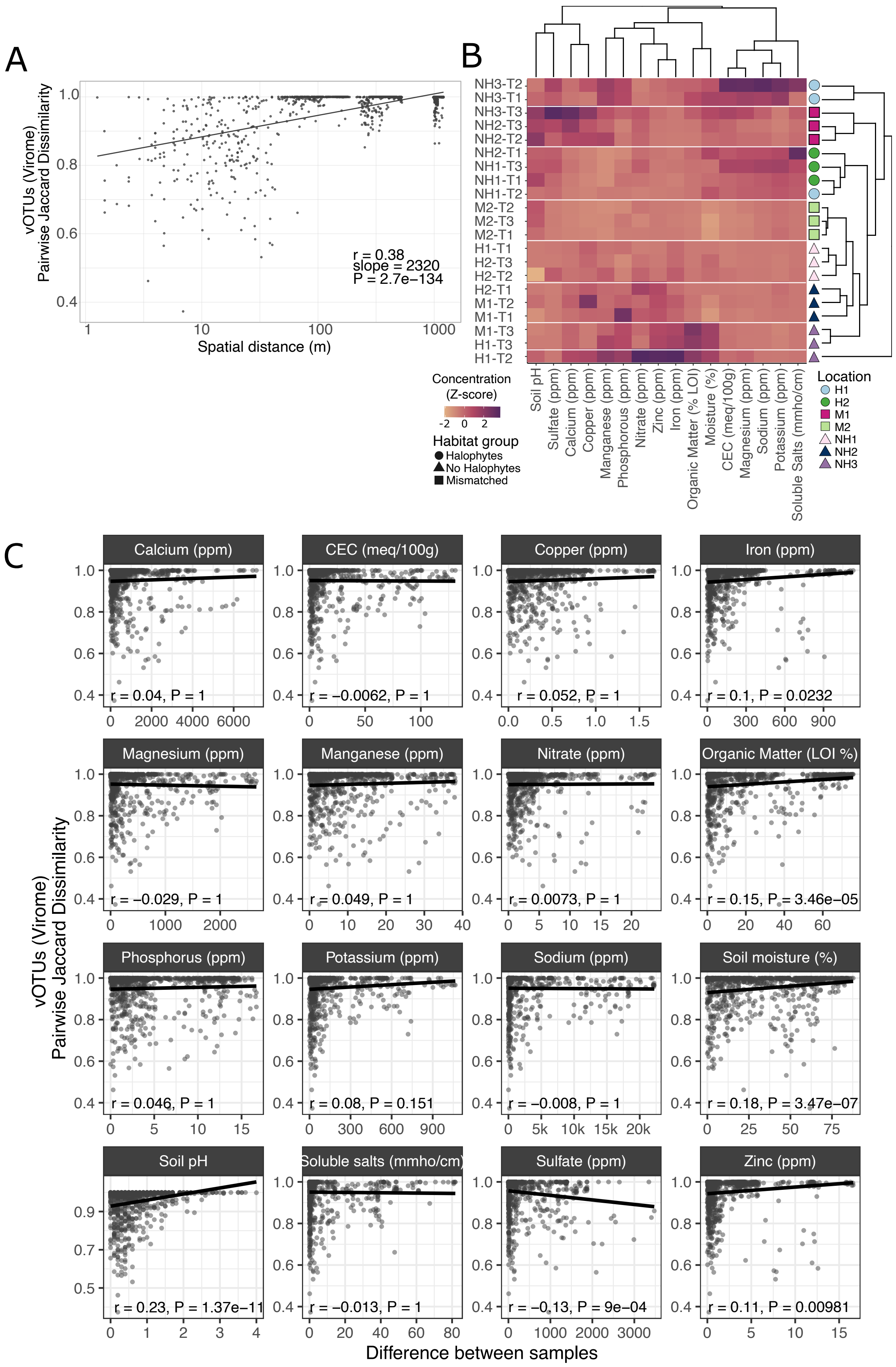


**Supplementary figure 2: A)** Relationship between Jaccard dissimilarity of viral communities and spatial distance (each point represents one pair of samples). **B)** Hierarchical clustering of average soil chemistry measurements across the three samples per site for each time point. The heatmap shows the average z-transformed value for each measurement. Shapes on the left indicate the wetland site. **C)** Relationship between Jaccard dissimilarity of viral communities and soil chemical measurements. ppm = parts per million, LOI = loss on ignition, CEC = cation exchange capacity.

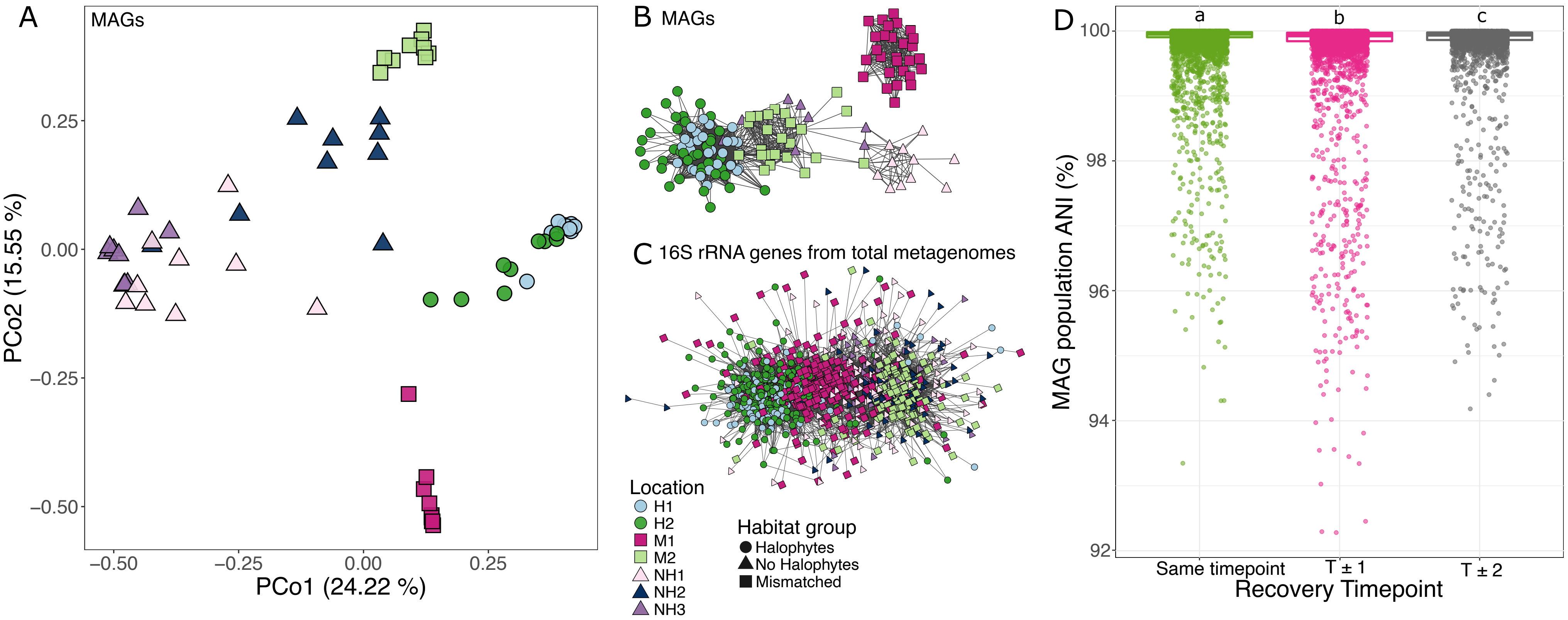


**Supplementary figure 3**: Prokaryotic community patterns corresponding to those explored for viral communities in the main text. **A)** Principal coordinates analysis, based on Bray-Curtis dissimilarities of MAG community composition in 63 samples. **B,C)** Co-occurrence network of **B)** MAGs and **C)** 16S rRNA gene OTUs (from total metagenomes) recovered in more than one sample. Each node represents a MAG or OTU, colored by the site in which it was most abundant (in which average per bp coverage was highest), and each edge represents significant co-occurrence between two nodes, calculated using a probabilistic co-occurrence model **D)** Prokaryotic population (MAG variant) microdiversity at Bodega Bay in 2021. Pairwise average nucleotide identities (ANIs) between MAG variants are plotted, calculated between each sample-specific MAG consensus sequence and the originally assembled (reference) MAG sequence, using inStrain. Each point is the ANI for one MAG variant compared to the reference sequence for that MAG. Box plots represent MAGs either assembled and recovered at the same time point, or at different timepoints, where T±1 equals 2 months between samplings, and T±2 equals 4 months. Letters above boxes correspond to significant differences between groups (Student's T test, significant when $p < 0.01$. Boxes show the median and interquartile range (IQR), and whiskers extend to Q1-1.5*IQR and Q3+1.5*IQR.

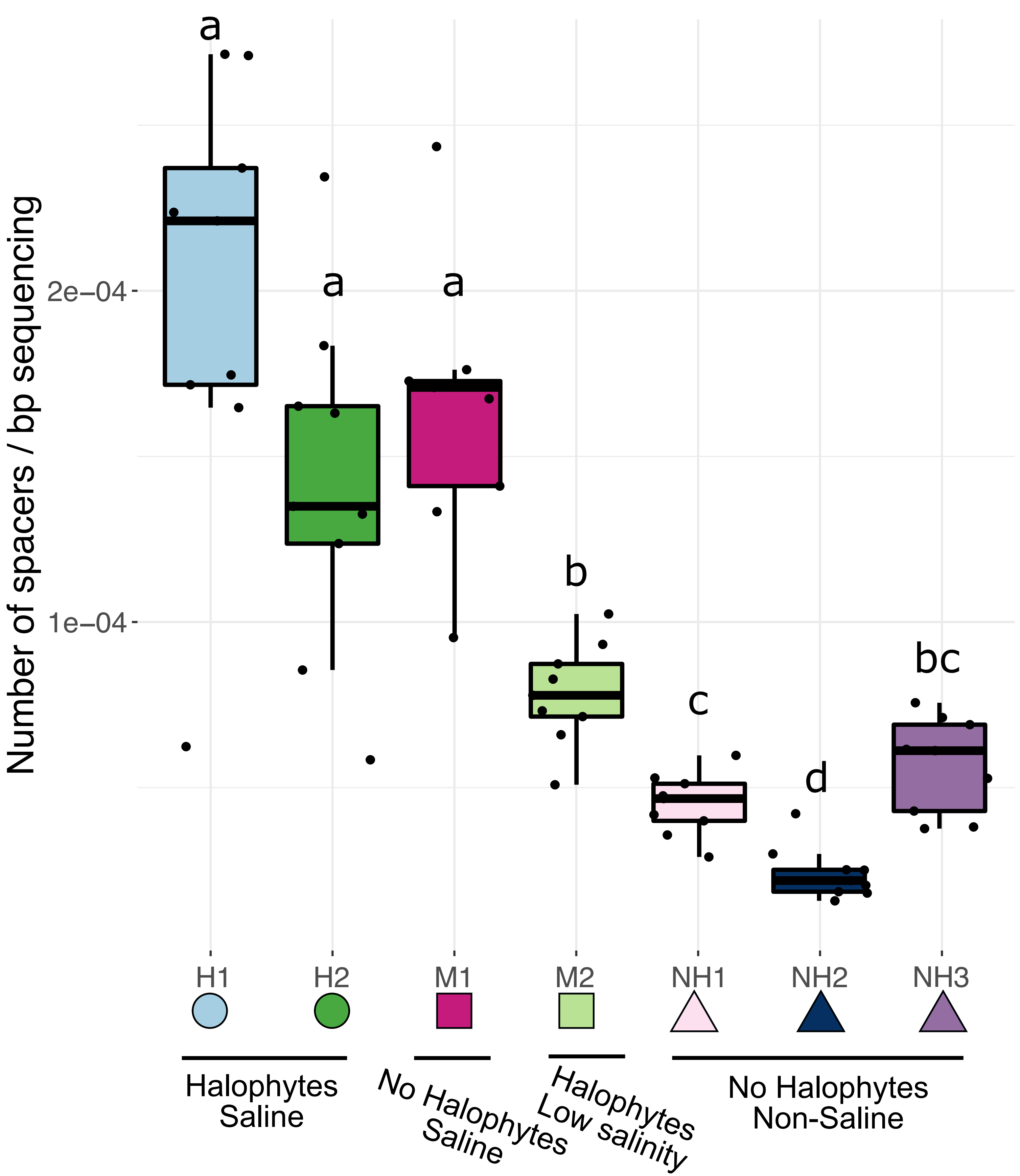


**Supplementary figure 4:** Number of predicted CRISPR-spacer arrays per bp of metagenomic sequence for each of the seven wetland sites.

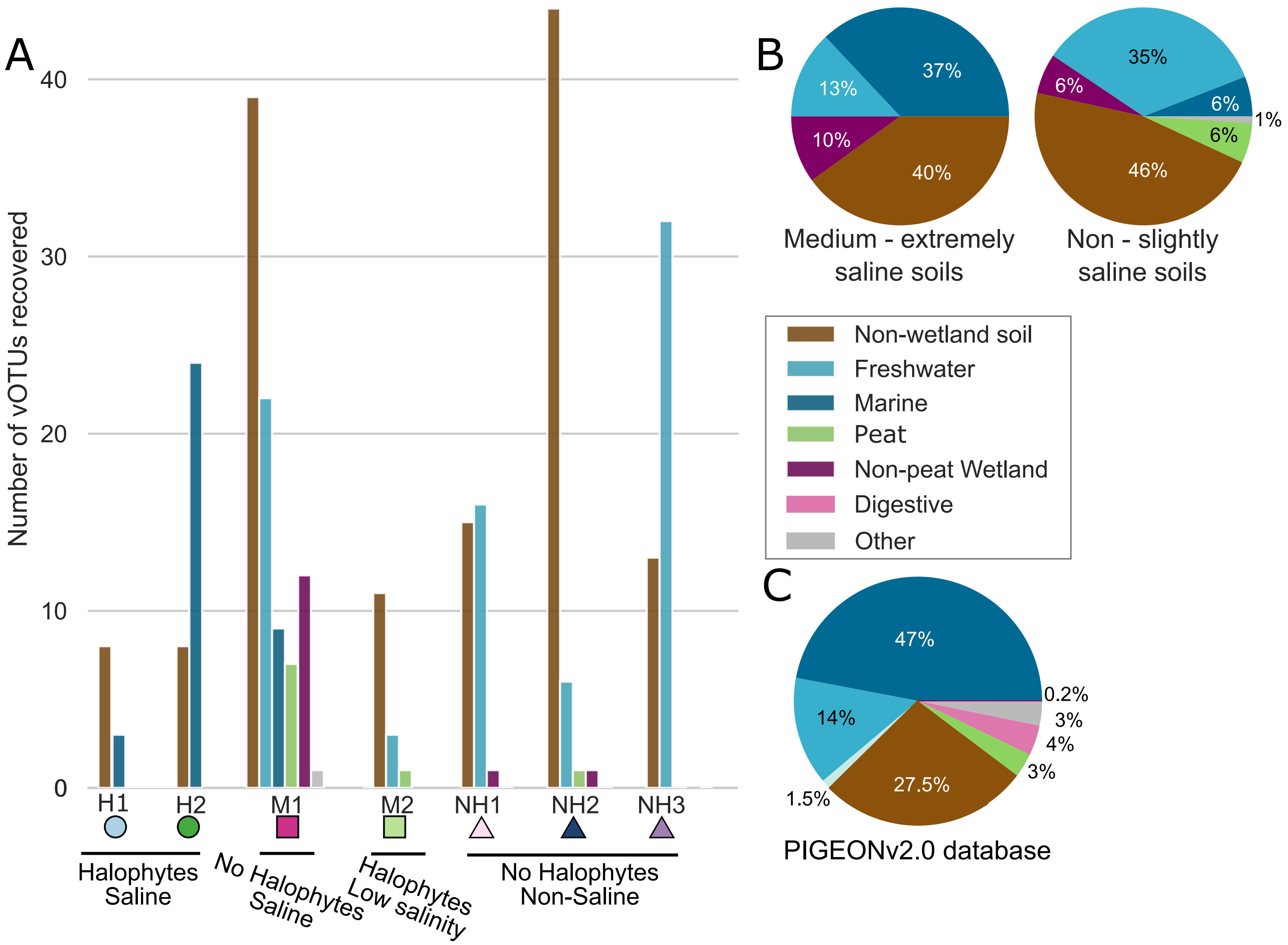


**Supplementary figure 5: A)** Total number of vOTUs recovered via viromic read mapping against PIGEONv2.0 per wetland site, colored by original source environment in PIGEONv2.0. **B)** Relative proportions of vOTUs recovered from PIGEONv2.0 at Bodega Bay, according to soil salinity group for their Bodega Bay site, colored by original source environment from PIGEONv2.0. **C)** Composition of the PIGEON v2.0 database of 515,763 vOTU sequences (same as in Figure 2B, repeated here for ease of comparison to panels A and B).
